# Publicly available hiPSC lines with extreme polygenic risk scores for modeling schizophrenia

**DOI:** 10.1101/2020.07.04.185348

**Authors:** Kristina Rehbach, Hanwen Zhang, Debamitra Das, Sara Abdollahi, Tim Prorok, Sulagna Ghosh, Sarah Weintraub, Giulio Genovese, Samuel Powell, Anina Lund, Schahram Akbarian, Kevin Eggan, Steven McCarroll, Jubao Duan, Dimitrios Avramopoulos, Kristen J. Brennand

## Abstract

Schizophrenia (SZ) is a common and debilitating psychiatric disorder with limited effective treatment options. Although highly heritable, risk for this polygenic disorder depends on the complex interplay of hundreds of common and rare variants. Translating the growing list of genetic loci significantly associated with disease into medically actionable information remains an important challenge. Thus, establishing platforms with which to validate the impact of risk variants in cell-type-specific and donor-dependent contexts is critical. Towards this, we selected and characterize a collection of twelve human induced pluripotent stem cell (hiPSC) lines derived from control donors with extremely low and high SZ polygenic risk scores (PRS). These hiPSC lines are publicly available at the California Institute for Regenerative Medicine (CIRM). The suitability of these extreme PRS hiPSCs for CRISPR-based isogenic comparisons of neurons and glia was evaluated across three independent laboratories, identifying 9 out of 12 meeting our criteria. We report a standardized resource of publicly available hiPSCs, with which we collectively commit to conducting future CRISPR-engineering, in order to facilitate comparison and integration of functional validation studies across the field of psychiatric genetics.

## INTRODUCTION

Schizophrenia (SZ) is a disorder marked by extremely heterogeneous clinical presentation and an equally complex genetic risk architecture, with contributions from common and rare variants (reviewed ^1^). To date, genetic studies have identified 145 loci with common single nucleotide polymorphisms (SNPs) of small effect sizes ^2^, eight rare copy number variations (CNVs) of relative high penetrance ^3^, and two genes with rare but highly penetrance loos-of-function mutations (*SETD1A* ^4^; *RBM12* ^5^) that are significantly associated with risk for SZ. Whereas rare variants are typically highly penetrant, common variants individually result in small increases in the odds ratio (OR < 1.3) for SZ ^6^, accounting for a substantial portion of disease risk only in aggregate. There is a growing consensus that genetic risk converges between psychiatric ^7^, but not neurodegenerative disorders^8^, particularly focused on genes expressed during fetal cortical development ^9–12^, and involved in synaptic biology ^3, 13–16^ or gene regulation ^13, 14, 17^.

Although genome wide association studies (GWAS) ^2^ and large scale exome sequencing projects (https://schema.org/) are increasingly dissecting the genetic architecture of risk for psychiatric disease, clinical translation of this knowledge are lagging. Many disease-associated variants are noncoding, suggesting a role in gene regulation ^18^; nearly half of the known SZ SNPs likely differentially regulate nearby gene expression ^19^. Although the regulatory impact of these SNPs can be predicted by overlapping GWAS with multi-omic transcriptomic (expression quantitative trait loci, eQTL ^20, 21^) and epigenetic (histone modification ^22^, and chromatin accessibility ^23^ and looping ^24, 25^) data sets, functionally demonstrating how these loci act as causal contributors to disease risk remains an intractable problem. There is an increasing need to functionally dissect the causal mechanisms underlying disease risk ^26, 27^, as many important questions remain unanswered. What are the mechanisms by which this growing list of common single nucleotide polymorphisms (SNPs) act within the diverse cell types of the brain to contribute to SZ etiology? Is there pathway level convergence of these disorder-associated genes, and if so, does convergence occur in a cell-type-specific manner? Do these common variants of small effect interact in a strictly additive fashion^26^, or through more complex epistatic ^28^ or omnigenic models ^29^? Systematic functional validation of these many risk variants requires a more tractable and amendable model. For instance, one that can perform network level perturbations to understand GWAS risk variants that may act additively and/or synergistically ^30^.

Theoretically capable of generating every cell type in the human body, human induced pluripotent stem cells (hiPSCs) are a useful tool with which to generate patient-specific cells for functional genomic studies. hiPSC-derived neurons and glia most resemble their prenatal in vivo counterparts ^31–36^, and are particularly well-suited to test the neurodevelopmental impact of the many psychiatric risk variants predicted to exert their influence during fetal cortical development ^12^. Emerging brain organoids ^37^ and more complex assembloid ^38–40^ models enable studies with extended maturation periods in more physiologically relevant contexts. Although case/control hiPSC-based models demonstrate concordance with post-mortem data sets ^36^, common genetic differences between donors (independent of diagnosis) leads to substantial inter-individual heterogeneity between hiPSC lines ^41, 42^. Given the limited sample sizes feasible in hiPSC studies, the most rigorous and well-powered approach to study individual variants is through isogenic comparisons on the same donor background ^43^. The isogenic approach has been recently made possible by the CRISPR-based genome editing toolbox ^44^ to empirically evaluate the impact of SZ-associated changes in DNA sequence, endogenous gene expression, DNA methylation, histone modification and chromatin confirmation (reviewed ^45^). We ^23, 46, 47^ and others ^48, 49^ have applied genome editing to facilitate the empirical validation of putative causal variants through precise isogenic comparisons on defined genetic backgrounds. However, due to the limited number of hiPSC lines used in each study while the cellular phenotypic expressivity may be influenced by genetic backgrounds, we are so far unable to evaluate and compare the effect. Thus, the field is in need of a set of hiPSC lines with defined genetic backgrounds that can be used by different investigators to produce data that can be cross-replicated and integrated.

Towards the goal of facilitating functional genomic studies of how risk variants interact with donor background in a cell-type specific manner, here we characterized the genomic integrity, pluripotency, and gene transfection and neural differentiation efficiencies, as well as CRISPR genome edibility of twelve control hiPSCs derived from donors with exceptionally high and low polygenic risk scores (PRS) for SZ **(Fig. 1)**. Through complementary studies conducted at all three sites, we collectively identified the nine hiPSCs most appropriate for neuronal induction and CRISPR-editing. These publicly available hiPSC lines with extreme SZ PRS will not only help evaluate the reproducibility of functional genomic studies, but also facilitate the integration of datasets generated by different laboratories, revealing any convergent impacts of independent SZ-GWAS genes/variants.

**Figure 1.**
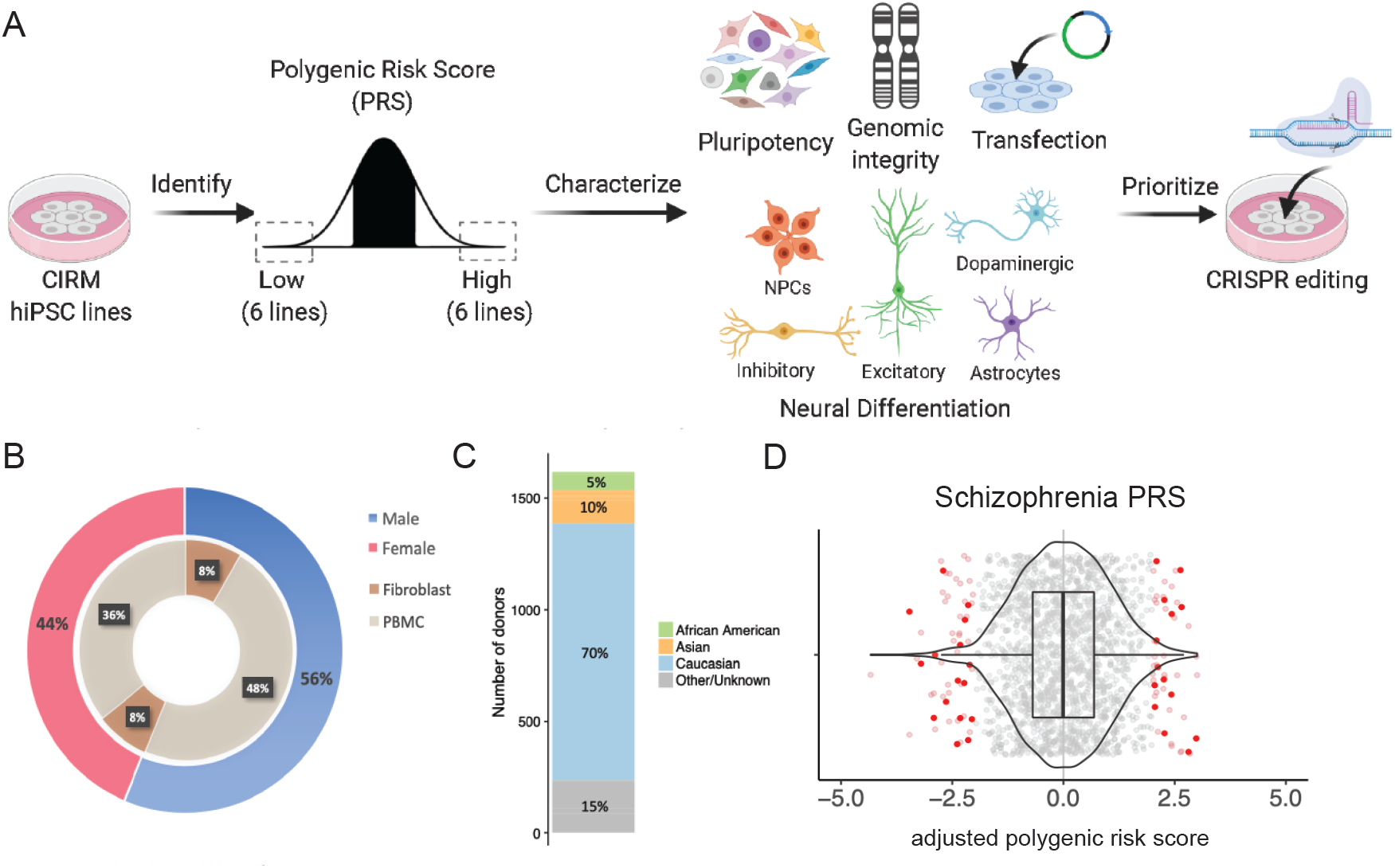
Identification, characterization and prioritization of high and low schizophrenia PRS hiPSC lines. **A.** Schematic summary of analysis pipeline. Figure created with BioRender.com. **B.** Overview of the 1628 hiPSC lines in the CIRM repository, showing gender and source of hiPSCs. **C.** Reported Ethnicities of CIRM hiPSC lines. **D.** Identification of hiPSC lines with >2 standard deviations (SD) lower or higher polygenic risk score (PRS) for schizophrenia based on PGC2.

## RESULTS

### Identification of hiPSCs derived from donors with extreme high and low PRS for SZ

The California Institute of Regenerative Medicine (CIRM) hiPSC collection is one of the largest USA-based single-deriver collections of genotyped hiPSCs (1618 donors); generated by a standardized, non-integrating episomal reprogramming approach in a single production facility (http://catalog.coriell.org/CIRM). Genome-wide SNP genotype data (from SNP microarray Illumina Infinium HumanCore BeadChip) was used to derive the SZ PRS scores using Psychiatric Genomics Consortium (PGC) 2 GWAS summary statistics ^2^ for all the non-SZ donors. We identified high and low PRS hiPSC lines with an adjusted PRS >2 standard deviations, and therefore with an expected >10-fold relative risk for disease ^2^. After excluding hiPSCs with potential sample swaps, gender discrepancies and structural variants, this totaled 28 hiPSCs derived from peripheral blood mononuclear cells (PBMCs) from unrelated donors of European descent. From these, we prioritized hiPSCs from twelve unrelated high (three male, three female) and low (three male, three female) PRS donors (shaded, **Table 1**) for a collective tri-site evaluation (Icahn School of Medicine at Mount Sinai, University of Chicago, and Johns Hopkins University) to assess their suitability for neural induction and CRISPR-editing.

**Table 1.** Selection of high and low schizophrenia PRS hiPSC lines.

| DONOR ID | SEX | DESCRIPTION | AGE<br>(AT BIOPSY) | POLYGENIC<br>RISK<br>SCORE |
| --- | --- | --- | --- | --- |
| **CW30274** | **M** | **no psychiatric diagnosis** | **63** | **2.98** |
| **CW30525** | **F** | **no psychiatric diagnosis** | **56** | **2.81** |
| **CW20184** | **M** | **autism spectrum disorder (ASD)** | **24** | **2.65** |
| **CW30421** | **F** | **no psychiatric diagnosis** | **66** | **2.63** |
| **CW40201** | **F** | **no psychiatric diagnosis** | **76** | **2.43** |
| **CW30454** | **M** | **no psychiatric diagnosis** | **67** | **2.42** |
| CW20041 | F | no psychiatric diagnosis | 9 | 2.27 |
| CW30265 | F | no psychiatric diagnosis | 55 | 2.26 |
| CW20195 | F | autism spectrum disorder (ASD) | 24 | 2.12 |
| CW30350 | F | no psychiatric diagnosis | 12 | 2.09 |
| CW60130 | F | no psychiatric diagnosis | 6 | 2.09 |
| CW70016 | F | no psychiatric diagnosis | 49 | 2.06 |
| CW70191 | F | no psychiatric diagnosis | 61 | 2.05 |
| CW50094 | M | no psychiatric diagnosis | 70 | -2.06 |
| CW50106 | F | no psychiatric diagnosis | 67 | -2.11 |
| CW40187 | F | no psychiatric diagnosis | 71 | -2.14 |
| CW30484 | F | no psychiatric diagnosis | 23 | -2.14 |
| CW70004 | M | no psychiatric diagnosis | 65 | -2.21 |
| CW70280 | M | no psychiatric diagnosis | 59 | -2.23 |
| CW70164 | F | no psychiatric diagnosis | 83 | -2.32 |
| **CW30108** | **M** | **no psychiatric diagnosis** | **53** | **-2.32** |
| **CW70372** | **M** | **no psychiatric diagnosis** | **54** | **-2.37** |
| CW50101 | F | no psychiatric diagnosis | 78 | -2.39 |
| CW40067 | F | no psychiatric diagnosis | 76 | -2.70 |
| **CW70179** | **F** | **no psychiatric diagnosis** | **86** | **-2.88** |
| **CW30190** | **M** | **no psychiatric diagnosis** | **63** | **-2.91** |
| **CW70142** | **F** | **no psychiatric diagnosis** | **82** | **-3.20** |
| **CW30154** | **F** | **no psychiatric diagnosis** | **66** | **-3.46** |

### Validation of pluripotency and genomic integrity of extreme PRS hiPSCs

hiPSCs were rigorously validated by the CIRM IPSC repository for genomic integrity, pluripotency using a 48-gene classifier, and loss of the reprogramming transgenes (see **Methods**). All 12 extreme PRS hiPSC lines were subsequently re-evaluated by our three laboratories to confirm pluripotency and genomic integrity.

Two hiPSC lines demonstrated abnormal growth at two or more sites: low_PRS_4 showed low post-thaw viability and attachment, and high_PRS_3 showed low adhesion and spontaneous differentiation (**Table 2**). We confirmed pluripotency marker expression by immunohistochemistry (NANOG and TRA-1-60, **Fig. 2A**). The differentiation potential was evaluated by rapid five-day tri-lineage directed differentiation ^50^, following by qPCR against early lineage specific transcripts (ectoderm: *PAX6, LMX1A*; mesoderm: *HOPX*; ectoderm: *FOXP2*, *SOX17*; pluripotency: *NANOG*) (**Fig. 2B**).

**Table 2.**
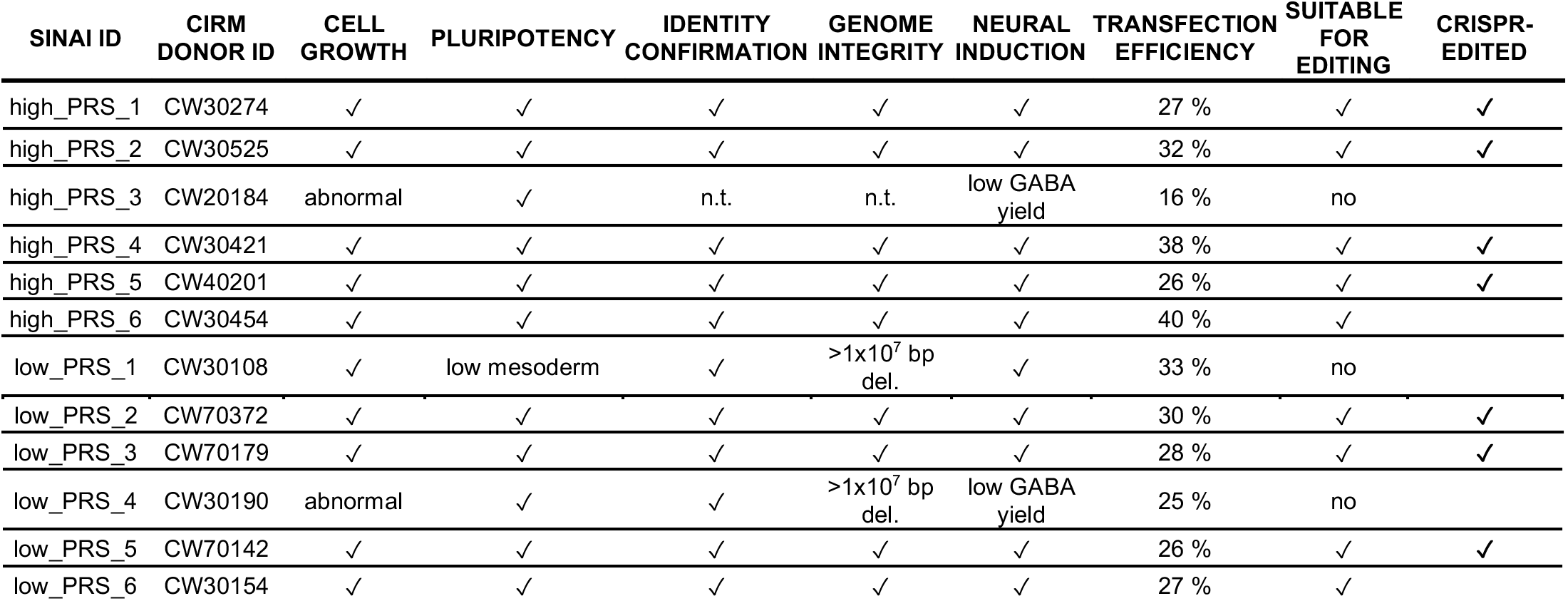
Summary of pluripotency, identity confirmation, genomic aberrations, neural induction, transfection efficiency and CRISPR-editing across 12 high and low PRS hiPSCs. (not tested, n.t.: high_PRS_3 was not fully evaluated owing to growth abnormalities.)

| SINAI ID | CIRM DONOR ID | CELL GROWTH | PLURIPOTENCY | IDENTITY CONFIRMATION | GENOME INTEGRITY | NEURAL INDUCTION | TRANSFECTION EFFICIENCY | SUITABLE FOR EDITING | CRISPR-EDITED |
| --- | --- | --- | --- | --- | --- | --- | --- | --- | --- |
| high_PRS_1 | CW30274 | ✓ | ✓ | ✓ | ✓ | ✓ | 27 % | ✓ | ✓ |
| high_PRS_2 | CW30525 | ✓ | ✓ | ✓ | ✓ | ✓ | 32 % | ✓ | ✓ |
| high_PRS_3 | CW20184 | abnormal | ✓ | n.t. | n.t. | low GABA yield | 16 % | no |  |
| high_PRS_4 | CW30421 | ✓ | ✓ | ✓ | ✓ | ✓ | 38 % | ✓ | ✓ |
| high_PRS_5 | CW40201 | ✓ | ✓ | ✓ | ✓ | ✓ | 26 % | ✓ | ✓ |
| high_PRS_6 | CW30454 | ✓ | ✓ | ✓ | ✓ | ✓ | 40 % | ✓ |  |
| low_PRS_1 | CW30108 | ✓ | low mesoderm | ✓ | >1x10<sup>7</sup> bp del. | ✓ | 33 % | no |  |
| low_PRS_2 | CW70372 | ✓ | ✓ | ✓ | ✓ | ✓ | 30 % | ✓ | ✓ |
| low_PRS_3 | CW70179 | ✓ | ✓ | ✓ | ✓ | ✓ | 28 % | ✓ | ✓ |
| low_PRS_4 | CW30190 | abnormal | ✓ | ✓ | >1x10<sup>7</sup> bp del. | low GABA yield | 25 % | no |  |
| low_PRS_5 | CW70142 | ✓ | ✓ | ✓ | ✓ | ✓ | 26 % | ✓ | ✓ |
| low_PRS_6 | CW30154 | ✓ | ✓ | ✓ | ✓ | ✓ | 27 % | ✓ |  |

**Figure 2.**
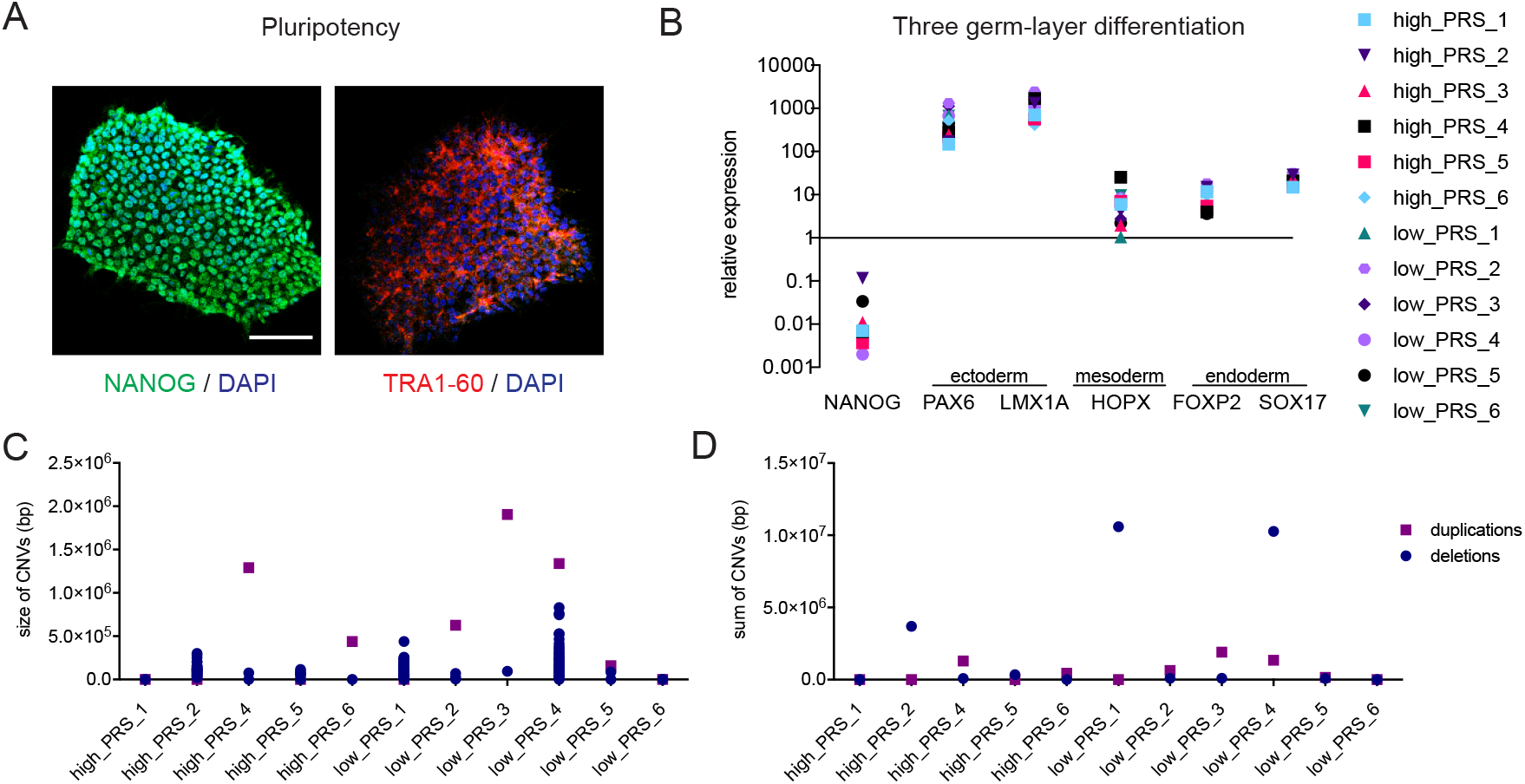
Validation, identity confirmation and CNV analysis of extreme PRS hiPSC lines. **A.** Representative immunofluorescence staining against the pluripotent markers NANOG and TRA-1-60. Scale bar: 100μm. **B.** Directed three germ layer differentiation of extreme PRS hiPSC lines for 5 days with subsequent qPCR analysis against typical lineage markers, normalizes to hiPSC control and 18S. **C.** The selected extreme PRS hiPSC lines have >99% concordance with the original CIRM data, confirming their identity. **D.** Identification of CNVs depicted by size. **E**. Sum of deletions and duplications present in the selected hiPSC lines after expansion.

hiPSCs were subjected to genotyping on the Multi-EthnicGlobal_D1 chip (illumina Inc.) to confirm donor identity against genotype data provided by the CIRM repository. The GenomeStudio CNV Region Report Plug-in v2.1.2 evaluated genome integrity and confirmed the absence of gross karyotypic abnormalities: high_PRS_4, low_PRS_3 and low_PRS_4 showed duplications >1.0×10^6^ bp, but more critically, high_PRS_4 exhibited an additional number of deletions >5.0×10^5^ bp, adding up to >1.0×10^7^ bp (**Fig. 2C-E**).

Overall, 10 of 12 extreme PRS hiPSCs showed robust proliferation, demonstrated pluripotency and genome integrity indicating overall suitability for functional genomic studies (**Table 2**).

### Demonstration of neuronal and glial generation

To ensure that the hiPSCs can robustly generate the neural subtypes most commonly linked to SZ (glutamatergic neurons ^51^, GABAergic neurons ^51^, dopaminergic neurons ^52^, and astrocytes ^53^), each laboratory independently evaluated neural differentiation and induction using different methodologies already well-established at each site ^23, 30, 54–56^.

Forebrain neural progenitor cells (NPCs) were generated from all 12 lines ^57^; immunofluorescence staining of PAX6 and NESTIN exceeded 95% in 10 out of 12 lines, with NPCs from two donors showing impurities (low_PRS_1 and low_PRS_4) (**Fig. 3A**). Glutamatergic neuron induction by overexpression of *NGN2* in hiPSCs ^58^ resulted in robust expression of glutamatergic genes by 21 days post-induction (**Fig. 3A**). GABAergic neuronal induction via overexpression of *DLX2* and *ASCL1* was efficient from both hiPSCs ^59^ and NPCs ^54^ (**Fig. 3A**). 32 days post-induction, NPC-induced GABAergic neurons expressed GABAergic neuronal genes glutamate decarboxylase 2 (*GAD2*) and vesicular GABA transporter (*vGAT*) by qPCR (**Fig. S1**). GABAergic neurons were stained with HuNu, MAP2 and GABA, demonstrating ~80% yield for 9 out of 12 lines (exceptions included 65% for low_PRS_5 and ~50% for high_PRS_3 and low_PRS_4) (**Fig. 3 A,B**). Those hiPSCs with reduced propensity for neuronal induction also had abnormalities in hiPSC culture and CNV analysis, suggesting that hiPSCs with irregular growth may be more likely to show defective differential potential as well. Dopaminergic neurons, a third neuronal cell type implicated in SZ, was achieved by a novel lentiviral induction strategy: overexpression of *ASCL1, NURR1* and *LMX1A* from hiPSCs ^60–62^. We derived dopaminergic neurons co-expressing tyrosine hydroxylase (TH) and synapsin (SYN) after 5 weeks of induction when co-culture with astrocytes for 6 of 12 hiPSC lines (**Fig. 3 A**); the variability between hiPSC lines and induction batches was high, so we recommend that dopaminergic neuron differentiation be individually optimized for the remaining hiPSC lines. We queried expression of the 67 non-MHC SZ risk genes associated by prediXcan ^63^ with the SZ-PGC2 loci ^2, 64^; finding that 62.6%, 79.1% and 76.1% are expressed at >0.5 log2RPKM, respectively, in our existing *NGN2*-glutamatergic, *ASCl1/DLX2*-GABAergic and *ASCL1, NURR1* and *LMX1A-* dopaminergic neuron RNAseq datasets ^54^.

**Figure 3.**
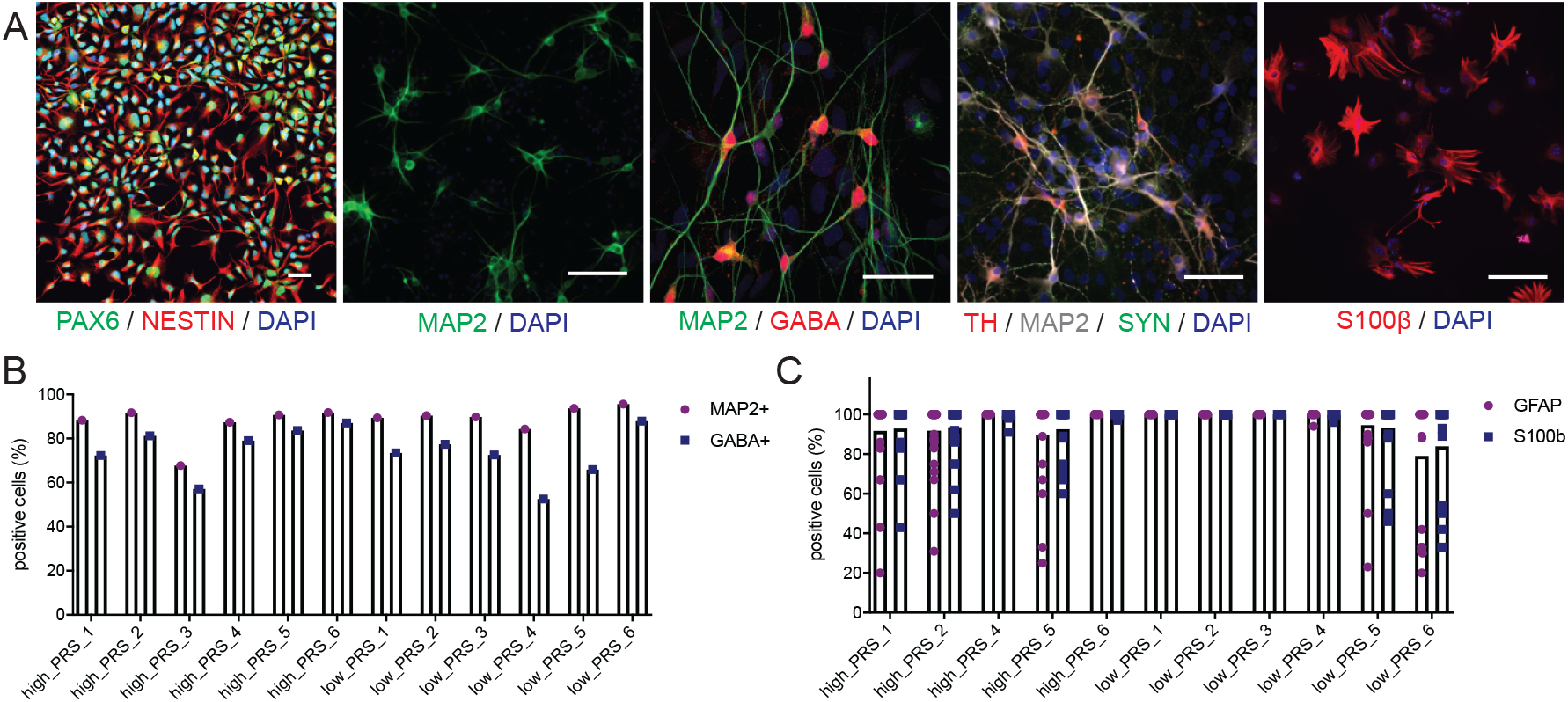
Demonstration of robust neuronal and glial generation. **A.** Representative images of astrocyte and neuron (glutamatergic, GABAergic, dopaminergic) differentiations. Scale bar: 50μm. **B.** Quantification of GFAP and S100b positive cells after 21 days of astrocyte differentiation. **C.** Quantification of MAP2 and GABA positive cells after 32 days of GABAergic neuron differentiation.

Lentiviral co-transduction of doxycycline-inducible *SOX9* and *NFIB* yields astrocytes ^65^. We have demonstrated high efficiency induction of astrocytes from the PRS hiPSCs, with 10 out of 11 hiPSCs yielding population of >90% GFAP and S100β positive cells by day 21, with only low_PRS_6 showing a slightly lower efficiency (~80%) (**Fig. 3 A,C**).

Overall, all 9 of 10 extreme hiPSCs with robust proliferation, demonstrated pluripotency and genome integrity showed strong capacity to generate neurons and glia through standard methods (**Table 2**).

### Transfection efficiency

An important perquisite for CRISPR editing is the ability to introduce Cas9, template DNA and gRNAs. hiPSC lines are commonly known to be difficult to genetically manipulate; optimization of transfection conditions and enriching for transfected cells are critical for efficiently recovering modified clones ^66^. To assess the likely efficiency of each hiPSC line for subsequent CRISPR engineering, we tested the efficiency of two commonly used methods to introduce DNA across our laboratories: transfection and nucleofection. We evaluated Lipofectamine STEM transfection and Amaxa nucleofection approaches with a GFP reporter followed by FACS analysis (**Fig. 4**). Lipofectamine methods yielded highly variable efficiencies that differed between sites (JHU: 6 to 73% (**Fig. 4 A**), UChicago: 16 to 23% (**Fig. 4 B**)). Nucleofection was generally more efficient: 36 to 65% (**Fig. 4 C**). Across all methods and sites, the two hiPSCs without lowest transfection efficiencies (high_PRS_3 and low_PRS_4) consistently showed the poorest culture and neuronal generation as well.

**Figure 4.**
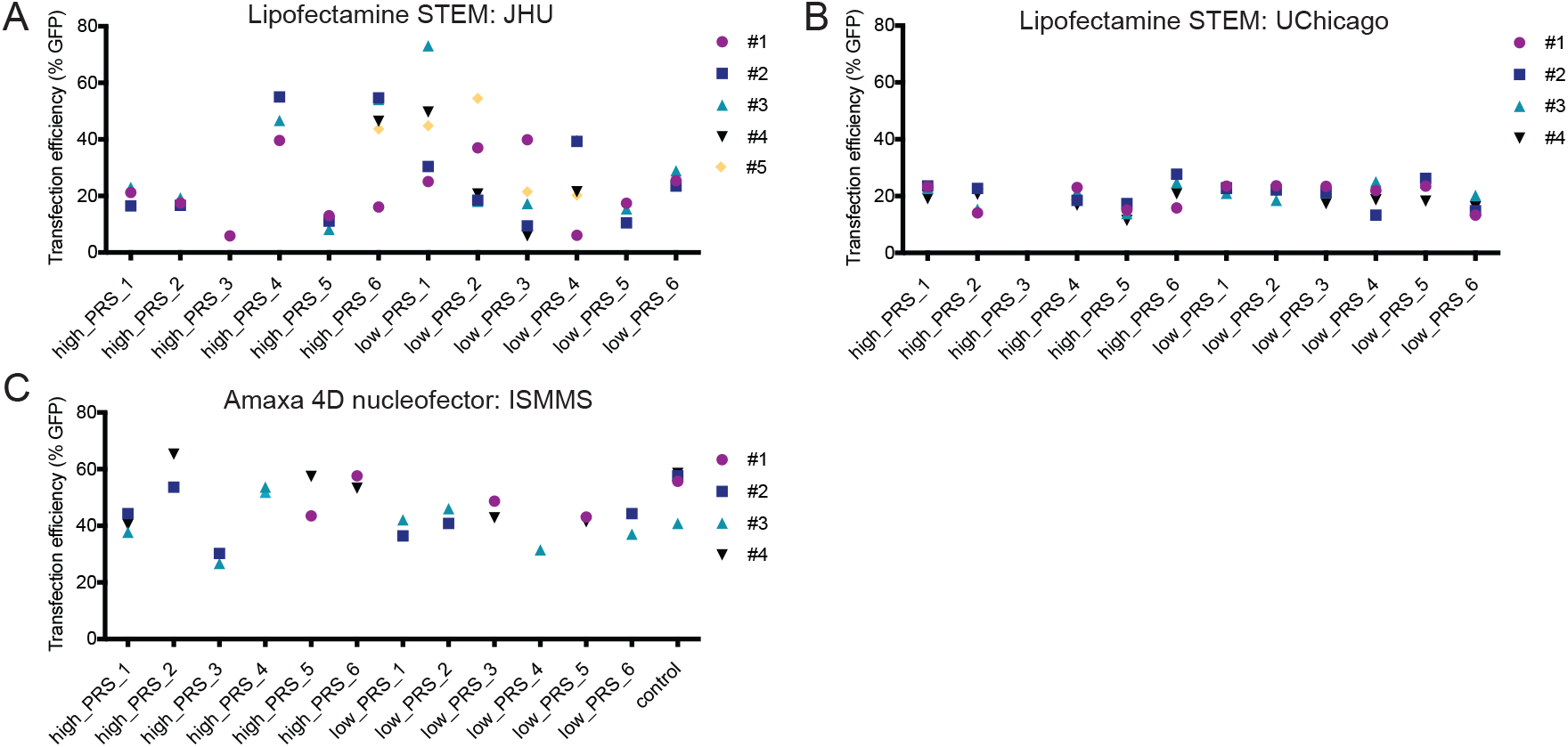
Appropriateness for CRISPR-editing. Transfection efficiencies using Lipofectamine STEM and the Lonza 4D nucleofector across three different sites: JHU, ISMMS, UC.

Overall, from 9 of 9 extreme hiPSCs with robust proliferation, demonstrated pluripotency, genome integrity, and neuronal/glia induction, we observed transfection efficiencies sufficient to achieve efficient CRISPR/Cas9 SNP editing (**Table 2**).

### Appropriateness for CRISPR-editing

To evaluate the suitability for CRISPR/Cas9 editing, we applied two independent methodologies (Cas9-based editing ^67^ and dCas9-based DNA base editing ^68^) against two independent non-coding SNPs already edited in our laboratories.

First, we re-targeted the SZ risk SNP rs4702 within the 3’ region of the gene *FURIN* ^46^. hiPSC lines were nucleofected with a Cas9 protein/gRNA complex and a respective repair oligonucleotide for either the wildtype allele A or the risk allele G (**Fig. 5 A,B**). Bulk analysis of editing efficiency was performed by restriction enzyme digest, revealing efficient editing in both directions (whether starting from either heterozygous allele (low_PRS_5, low_PRS_2, high_PRS_1) or homozygous AA alleles (high_PRS_2)). Correctly edited clones were identified from 5 of 5 hiPSC lines evaluated, with efficiencies of 21.7%, 25%, 52.2%, 50%, 27.8%, 26% and 37.5% respectively (**Fig. 5C**).

**Figure 5.**
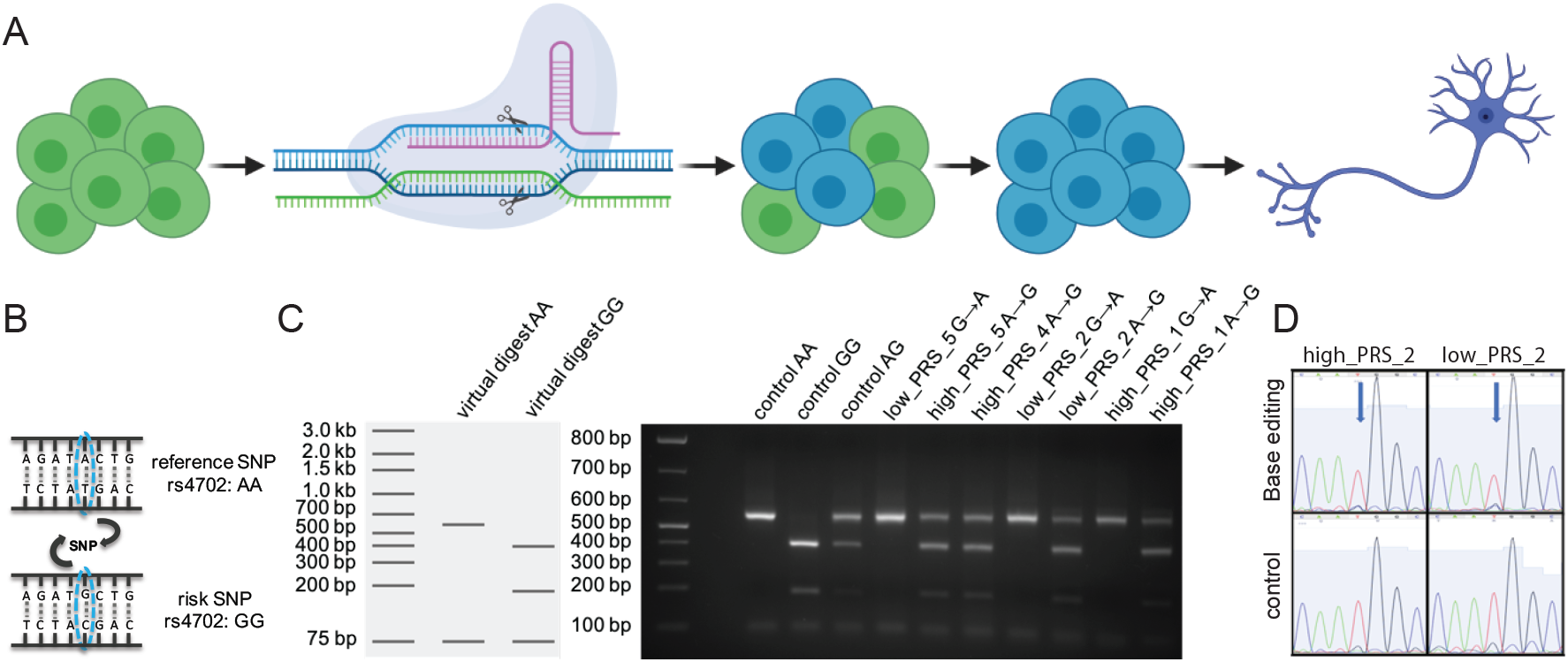
Demonstration of CRISPR-editing of schizophrenia risk SNPs. **A.** Overview of editing pipeline, using CRISPR/Cas9 and a repair oligonucleotide to achieve homologous recombination (HDR). Figure created with BioRender.com. **B.** Example of schizophrenia GWAS-SNP rs4702, with reference allele A and risk allele G. **C.** Bulk editing analysis of *FURIN* rs4702 using the restriction enzyme SfANI, cutting the A allele once and the G allele twice. **D.** Sanger sequencing results from base editing of two extreme PRS hiPSC lines. Top row: *BAG5* rs7148456 edited (T to C), with blue arrow pointing to SNP; bottom row: unedited homozygous controls. Note the small peak of the edited C allele that was absent in the unedited controls and a much lower peak of T allele in the edited sample vs. unedited controls.

Second, we applied dCas9 to re-target the SZ risk SNP rs7148456 associated with allele-specific open chromatin at the *BAG5* locus ^55^. In two hiPSC lines, we transfected a customized ABEmax vector system ^68^ to express the gRNA, dCas9 and ABEmax in a single vector. This DNA base editor is expected to have higher editing efficiency than homologous-directed repair in CRISPR/cas9 SNP editing, with minimal DNA off-target editing because the deactivated Cas9 does not create double-strand DNA breaks ^68^. hiPSCs were transfected using Lipofectamine STEM reagent. After puromycin screening, bulk DNAs were extracted approximately 72 hours post transfection and sequenced. We approximated the editing efficiencies (A to G) based on A to G (or T to C on the minus strand) percentage by Sanger sequencing, which showed a 11.2% and 13.8% editing efficiency for high_PRS_2 and low_PRS_2, respectively (**Fig. 5D**).

Overall, while we believe that all 9 validated extreme PRS hiPSC (5 high and 4 low) should be suitable for CRISPR-editing, for 7 we have empirically demonstrated efficient allelic conversion of a single non-coding SNP (**Table 2**).

## DISCUSSION

A major challenge for the field of psychiatric genetics is translating highly complex genetic insights into medically actionable information. The integration of hiPSCs and CRISPR-editing is becoming the preferred means to test the function of common and rare disease-associated variants. Many studies now follow a design whereby CRISPR-edited isogenic lines are contrasted; however, there is tremendous variability in the efficiency with which different hiPSC lines respond to CRISPR-editing and neuron differentiation ^69^. Towards this, we conducted a collaborative and systematic tri-site validation of the pluripotency, genomic integrity, propensity for neuronal differentiation and CRISPR-editing capabilities of six high and six low PRS control hiPSCs, prioritizing nine lines suitable for functional validation of SZ variants.

This publicly available resource of well-characterized extreme PRS hiPSC lines will facilitate the study of schizophrenia risk in a well-defined cohort that is comparable across different sites. High and low PRS lines can be used to address the question of penetrance when modeling rare or common variants, to examine the potential impact of genetic background on phenotypic expressivity in hiPSC derived cells. Using this well-characterized cohort, will reduce the time and resources spent validating CRISPR-editing conditions across hiPSCs of lesser quality or with less available genetic information. Eventually, if adopted across additional psychiatric genetics laboratories, we will improve the reproducibility of hiPSC-based studies between research groups and increase the power of isogenic CRISPR-based experiments by facilitating meta-analyses with an ever-growing dataset of isogenic hiPSCs. Most importantly, this resource will facilitate the future integration of datasets generated across laboratories. Such meta-analyses identify convergent effects between SZ-GWAS risk variants.

We note three limitations to this resource. First, while we expect that the concept of shared extreme PRS hiPSCs will prove to be of enduring value, this specific hiPSC cohort might have to be re-defined as PRS calculations improve (by expanded PGC3-GWAS cohort size ^70^ and/or incorporation of pathway-specific information ^71^), which could necessitate the inclusion of additional extreme PRS hiPSCs. Second, being derived entirely of donors of European descent, this cohort is not well suited to test trans-ancestry effects. Third, and most importantly, this collection is unlikely to prove to be adequately powered to resolve sex-specific and/or high versus low PRS effects, and is mostly intended for the validation of isogenic functional genomic studies across a diverse collection of donor backgrounds. Of course, experimentally deconvolving the extent to which individual risk factors exert variable effects across extreme PRS donor backgrounds will help to resolve the extent to which polygenic risk reflects additive^26, 72^ or more complex epistatic^28^ or omnigenic models ^29^ of inheritance, and will in turn shape future refinements of PRS.

In summary, we describe a publicly available hiPSC collection from donors with extreme PRS for SZ and characterized the suitability to test functional effects of SZ risk variants in a context-dependent manner. As a shared resource, these hiPSC lines will enable the cross-lab reproducibility evaluation and data integration. However, given the polygenic nature of SZ and other neuropsychiatric disorders, we acknowledge that a larger number of iPSC lines of extreme SZ PRS would be needed for delineating the effect of polygenic risk background on disease modeling. Nonetheless, we anticipate that a similar strategy will prove broadly useful across many complex genetic disorders, although of course each disorder will require identification and validation of a unique set of extreme PRS hiPSCs.

## METHODS

### Prioritization of high and low PRS hiPSCs

There is uniform demographic and clinical data for the entire the CIRM hiPSC cohort (https://www.cirm.ca.gov/researchers/ipsc-repository). All available lines were validated across the following metrics: 1) chromosomal integrity (Illumina Infinium HumanCore BeadChip), no amplifications larger than 5MB (resolution of traditional G-banding assay) on SNP arrays with LogRDev score less than 0.5 that were not pre-existing in the donor; 2) pluripotency (qPCR of 48 mRNAs), a non-probabilistic binary linear classifier identifies the gene expression of the sample as iPSC based on an appropriate training set; 3) identity confirmation (PCR assay for 48 SNPs), ≤ 1 mismatch between donor and hiPSC line; 4) loss of reprogramming transgenes (PCR for two plasmid EBNA and OriP sequences), detection of ≤ 1 plasmid copy per 100 cells or a decrease in the number of plasmid copies detected at passage five; 5) mycoplasma negative (qPCR for 8 species); 6) sterility (microbiological testing by third-party service provider). We selected only those hiPSCs derived from a common somatic cell source (peripheral blood mononuclear cells, PBMCs), with an absence of acquired structural variants and gender discrepancies, and from unrelated donors of European ancestry, as SZ PRS was calculated from European subjects ^2^ and is ancestry dependent ^73^. Out of 30 samples with a PRS of at least 2 standard deviations from the mean, 3 female and 3 male subjects were selected per risk group.

### hiPSC culture

Human induced pluripotent stem cells (hiPSCs) were thawed with mTeSR1 (STEMCELL, #85850) and 1 μM H1152 (Millipore, #555550), seeded on Matrigel (Corning, #354230) coated 6-well plates and subsequently adapted to further culture in StemFlex media (Gibco, #A3349401). hiPSCs at ~ 70% confluence (1.5×10^6^ cells / well of a 6-well plate) were incubated in EDTA (Life Technologies #15575-020) for 4 min at room temperature (RT), the EDTA was aspirated, the cells dissociated in fresh StemFlex media and seeded onto Matrigel-coated plates at 3-5×10^5^ cells/well. Media was replaced every other day for 4 days until the next passage. For freezing and shipping to partners, cells were dissociated with EDTA and subsequently frozen in Stemflex medium with 10% DMSO (Sigma, # D2650) at −80°C and transferred to liquid nitrogen for long-term storage. All hiPSCs were tested and are mycoplasma free.

### Three germ-layer differentiation

After thawing, cells were passaged 1-2 times in mTeSR1 media before differentiation into the three germ layers using STEMdiff™ Definitive Endoderm Kit (STEMCELL, #05110), STEMdiff™ Mesoderm Induction Medium (STEMCELL, #05220) and dual-SMAD^74^ inhibition for ectoderm differentiation. hiPSCs were dissociated with Accutase Cell Detachment Solution (Innovative Cell Technologies, # AT-104) for 10min at 37 °C and seeded on Matrigel-coated 24 well plates in mTeSR1 medium with 1 μM H1152 at a density of 2.1×10^5cells/cm^2^ for endoderm and ectoderm differentiation and 5×10^4cells/cm^2^ for mesoderm differentiation. Endoderm and mesoderm differentiations were performed according to the manufacturers protocol, whereas ectoderm was induced with neural induction medium: DMEM/F12 (Thermofisher, #10565018), 1% N-2 supplement (Thermofisher, #17502048), 10μM SB431542 (Tocris, #1614), 200nM LDN193189 (Tocris, # 6053) with daily medium changes. Cells were harvested on day 5 and RNA was prepared using Trizol isolation.

### Neuron and Glia Generation

#### i. NPC differentiation from hiPSCs

Our NPCs were generated using PSC Neural Induction Medium (Thermofisher) and passaged using Rosette selection reagent (STEMCELL# 05832). To estimate purity, we stained PAX6 (Biolegend# PRB-278P) and NESTIN (Sigma# MAB5326).

#### ii. NGN2-glutamatergic neuron induction from hiPSCs ^58^

On day −1 a 6-well plate was coated with Matrigel. On day 0, hiPSCs were dissociated with Accutase Cell Detachment Solution (Innovative Cell Technologies, # AT-104), counted and transduced with rtTA (Addgene 20342) and *NGN2* (Addgene 99378) lentiviruses in StemFlex media containing 10 μM Thiazovivin (Millipore, #S1459). They were subsequently seeded at 1×10^6^ cells/well in the prepared 6-well plate. On day 1, Medium was switched to non-viral induction medium (DMEM/F12 (Thermofisher, #10565018), 1% N-2 (Thermofisher, #17502048), 2% B27-RA (Thermofisher, #12587010)) and doxycycline (dox) was added to each well at a final concentration of 1 μg/ml. At day 2, transduced hiPSCs were treated with 500 μg/mL G418 (Thermofisher, #10131035). At day 4, medium was replaced including 1 μg/ml dox and 4 μM cytosine arabinoside (Ara-C) to reduce the proliferation of non-neuronal cells. On day 5, young neurons were dissociated with Accutase Cell Detachment Solution (Innovative Cell Technologies, # AT-104), counted and seeded at a density of 1×10^6^ per well of a Marigel-coated12-well plate. Medium was switched to Brainphys neuron medium (Brainphys (STEMCELL, # 05790), 1% N2, 2% B27-RA, 1 μg/ml Natural Mouse Laminin (Thermofisher, # 23017015), 10 ng/ml BDNF (R&D, #248), 10 ng/ml GDNF (R&D, #212), 500 μg/ml Dibutyryl cyclic-AMP (Sigma, #D0627), 200 nM L-ascorbic acid (Sigma, # A4403)). For seeding, 10 μM Thiazovivin (Millipore, #S1459), 500 μg/mL G418 and 4 μM Ara-C and 1 μg/ml dox were added. At day 6, medium was replaced with Brainphys neuron medium with 4 μM Ara-C and 1 μg/ml dox. Subsequently, 50% of the medium was replaced with fresh neuronal medium (lacking dox and Ara-C) once every other day until the neurons were fixed or harvested at day 21.

#### iii. ASCL1/DLX2-GABAergic neuron induction from hiPSCs ^59^ and NPCs ^54^

GABAergic neurons were generated by from NPC by overexpression of *Dlx2* and *Ascl1*. NPCs were seeded at a density of 5.0 × 10^5^ cells per well of a 24-well tissue plate. At day-1, NPCs were transduced with CMV-*rtTA* (Addgene ID: 19780), TetO-*Ascl1*-T2A-*Puro* (Addgene ID: 97329) and TetO-*Dlx2*-IRES-*Hygro* (Addgene ID: 97330), incubated for 15 minutes at 37°C and spinfected at 1,000G for one hour at room temperature. At day 0, media was replaced with NPC media containing 1 μg/mL dox (Sigma, #D9891). At day 1, NPC media was replaced with NPC media containing 1 μg/mL dox 1 μg/mL puromycin (Sigma, #P7255) and 250 μg/mL hygromycin (ThermoFisher, 10687010). On day-7, NPCs were introduced to neuronal media via half-media changes. Cells received 1μg/mL of dox until DIV 14. To prevent the proliferation of dividing mitotic progenitors, 50 nM cytosineβ-D-arabinofuranoside, also known as Ara-C, (Sigma, #C6645) was supplemented in the neuronal media. Neurons were harvested at day 32 for analysis.

#### iv. ASCL1/NURR1/LMX1A-dopaminergic neuron induction from hiPSCs, adapted and modified from ^60–62^

On day −1 a 6-well plate was coated with Matrigel. On day 0, hiPSCs were dissociated with Accutase Cell Detachment Solution (Innovative Cell Technologies, # AT-104), counted and transduced with rtTA (Addgene 20342), *ASCL1, NURR1* and *LMX1* (Addgene #97329) lentiviruses. hiPSCs were mixed in a conical tube in low volume StemFlex media containing 10 μM Thiazovivin (Millipore, #S1459) and subsequently seeded at 1×10^6^ cells/well in the prepared 6-well plate. At day 1, medium was switched to non-viral induction medium (DMEM/F12 (Thermofisher, #10565018), 1% N-2 (Thermofisher, #17502048), 2% B27-RA (Thermofisher, #12587010)) and 1 μg/ml dox. On day 2, medium was replaced with Induction Media with1 μg/ml dox and 1 μg/ml Puromycin. At day 3, medium was replaced with Induction Media with1 μg/ml dox and 2 μg/ml Puromycin. On day 5 medium was replaced with Induction Media with1 μg/ml dox and 1 μg/ml Puromycin and 2 μM Ara-C. On day 6, antibiotic selection was withdrawn. On day 7 young neurons were dissociated with Accutase Cell Detachment Solution (Innovative Cell Technologies, # AT-104) and seeded at a density of 0.57 mio/cm^2^ in Brainphys neuron medium (Brainphys (STEMCELL, # 05790), 1% N2, 2% B27-RA, 1 μg/ml Natural Mouse Laminin (Thermofisher, # 23017015), 10 ng/ml BDNF (R&D, #248), 10 ng/ml GDNF (R&D, #212), 500 μg/ml Dibutyryl cyclic-AMP (Sigma, #D0627), 200 nM L-ascorbic acid (Sigma, # A4403)). For accelerated maturation, part of the neurons were seeded on a layer of human astrocytes. In this case, the medium was supplemented with 2% FBS (Thermofisher, #10082147). Subsequently, 50% of the medium was replaced with fresh neuronal medium once every other day until the neurons were fixed or harvested on day 35. Dox and Ara-C were withdrawn on day 14.

#### v. NFIB/SOX9-astrocyte induction from hiPSCs ^65^

On day 0, hiPSCs were hiPSCs were dissociated with Accutase Cell Detachment Solution (Innovative Cell Technologies, # AT-104), counted, transduced with rtTA (Addgene 20342), and inducible *Nfib* (Addgene # 117271) and *Sox9* (Addgene # 117269) lentiviruses. hiPSCs and viruses were mixed in a conical tube in low volume StemFlex media containing 10 μM Thiazovivin (Millipore, #S1459) and subsequently seeded at 0.75×10^6^ cells/well in the prepared 6-well plate. Expression was induced on day 1 in StemFlex media with 2.5 μg/ml dox (continued throughout). On day 2, medium was switched to Expansion medium (DMEM/F-12, 10% FBS, 1% N2 supplement) with 1 μg/ml puromycin and 200 μg/ml hygromycin for selection. On day 4 to 6, Expansion medium was gradually switched to FGF medium (Neurobasal, 2% B27 supplement, 1% NEAA, 1% Glutamax, and 1% FBS, 8 ng/ml FGF2, 5 ng/ml CNTF, and 10 ng/ml BMP4, from Peprotech). On day 7, cells were dissociated with Accutase Cell Detachment Solution (Innovative Cell Technologies, # AT-104) and replated in Matrigel-coated wells. On day 8 and 9, FGF medium was gradually switched to maturation medium (1:1 DMEM/F-12 and Neurobasal, 1% N2, 1% sodium pyruvate, and 0.5% Glutamax; 5 mg/ml *N*-acetyl-cysteine, 500 mg/ml dbcAMP; 5 ng/ml heparin-binding EGF-like growth factor, 10 ng/ml CNTF (Peprotech, #450-13), 10 ng/ml BMP4 (Peprotech, #120-05)). Subsequently, 50% of the medium was replaced with Maturation medium every 2–3 days.

### hiPSC CRISPR transfection and validation

#### i. hiPSC transfection JHU

Firstly, hiPSCs were seeded at a concentration of 250,000 cells/well of a six-well plate and when 30% confluent, typically on the third day, subject to lipofectamine transfection. The transfection medium was prepared by mixing two solutions. The first consisted of 2 μl/well pmaxGFP DNA (LONZA,1 μg/ul) diluted in 250 μl/well Opti-MEM® medium (ThermoFisher Scientific) to obtain 500 ng DNA/well concentration. The second is 10 μl/well LipofectamineTM STEM Transfection Reagent (ThermoFisher Scientific) diluted in 250 μl/well Opti-MEM® medium. The transfection medium consists of a 1:1 ratio of the first (pmaxGFP DNA and Opti-MEM®) and second (LipofectamineTM and Opti-MEM®) solutions that were incubated for 10 min before addition to the cells. Meanwhile, the cell medium was removed and replaced with 2 ml/well mixture of ROCK inhibitor and StemFlexTM medium (ThermoFisher Scientific) at 1:1000 ratio. After incubation, add 500 μl/well transfection medium to the cells with the ROCK inhibitor and StemFlexTM, then incubate at 37 °C for 4 hrs. Afterward, add 2 ml/well of StemFlexTM medium to the mix and incubate at 37 °C overnight. After about 24 h, change the media and replace with 2 ml/well of StemFlexTM medium, then incubate at 37 °C overnight. After about 48 hrs, the cells can be washed, lifted and analyzed by fluorescence-activated cell sorting (FACS). Note that controls were prepared following a similar procedure, but instead of pmaxGFP DNA, the same volume of water was added.

#### ii. hiPSC transfection UC

All iPSC lines were passaged according to manufacturer’s protocols. Briefly, 50,000 cells were plated on Matrigel (Corning #354234) coated 24-well plates with mTeSR1 (Stemcell, #85850) media. Twenty-four hours after plating cells each well is treated with 1.2 μl Lipofectamine STEM reagent (ThermoFisher, #STEM00001) and 500 ng pmaxGFP (Lonza, PBP3-00675) combined into 50μl Opti-MEM media following Lipofectamine STEM reagent protocol. The cells were overlaid with fresh mTeSR1 the following day. Forty-eight hours post transfection the cells are washed with 1× PBS, treated with Accutase (Stemcell, cat# 07922) for 10 mins followed by 200 g centrifugation. The supernatant is removed and cells resuspended in fresh mTeSR media. The cell suspension is pipetted through Falcon tubes with cell-strainer caps and then analyzed by MACSQuant Analyzer 10 Flow Cytometer (Miltenyi Biotec) for 10,000 events.

#### iii. hiPSC nucleofection IMSSM

hiPSCs were transfected using the Lonza P3 Primary Cell 4D-Nucleofector Kit (V4XP-3024) according to the manufacturer’s instructions. Briefly, cells were dissociated following 15 min Accutase Cell Detachment Solution (Innovative Cell Technologies, # AT-104) incubation at 37°C and 1×10^6^ cells were centrifuged at 800 × g for 5 min. Pellet was resuspended in 100 μl nucleofector solution containing 800 ng GFP plasmid. The suspension was quickly transferred to a nucleofection cuvette, transfected in the Lonza 4D nucleofector program ‘CA-137’and seeded onto a Matrigel-covered 6-well plate in StemFlex containing 1 μM H1152 (Millipore, #555550). On the next day, medium was replaced. Two days after nucleofection, cells were dissociated using Accutase and cells were stained with LIVE/DEAD™ Fixable Far Red Dead Cell Stain Kit (Thermofisher, #L34973). The cells were washed with PBS, filtered using FACS tubes with cell-strainer caps and subsequently analyzed using the BD FACSCanto™ ‖ counting 10,000 events.

#### iv. hiPSC SNP rs4702 editing using Cas9 protein

hiPSCs were transfected with 10 μg TrueCut™ Cas9 Protein v2 (ThermoFisher, # A36498), 8 μg synthetic gRNAs (Synthego): FURIN rs4702 A-G conversion: GGCTGGTTTTGTAAGATACT; FURIN rs4702 G-A conversion: GGCTGGTTTT-GTAAGATGCT and 100pmol of the respective repair ssODNs (ThermoFisher): rs4702 G->A: TZFFATAGAACCAGCAATGCTGGGCCTGTTTAAATTACAAGAAAAAAATCACT GTGCACCAACCCAGZATCTTACAAAACCAGCCGGGCTGGCCAOEA; rs4702 A->G: TZFFATAGAACCAGCAATGCTGGGCCTGTTTAAATTACAAGAAAAAAATCACT GTGCACCAACCCAGOATCTTACAAAACCAGCCGGGCTGGCCAOEA using the Lonza P3 Primary Cell 4D-Nucleofector Kit (V4XP-3024). Briefly, cells were dissociated following 15 min Accutase Cell Detachment Solution (Innovative Cell Technologies, # AT-104) incubation at 37 °C and 1×10^6^ cells were centrifuged at 800 × g for 5 min. Pellet was resuspended in 100 μl nucleofector solution containing the assembled. The suspension was quickly transferred to a nucleofection cuvette, transfected in the Lonza 4D nucleofector program ‘CA-137’and seeded onto a Matrigel-covered 6-well plate in 3 ml StemFlex containing 10 μM Thiazovivin (Millipore, #S1459). The cells were subjected to a 48 hour cold shock at 32 °C to enhance homology directed repair (HDR). One day after nucleofection, the medium was replaced by fresh StemFlex medium without Thiazovivin. The medium was changed every other day, until 70-80% confluence. Cells were dissociated with Accutase, the majority of cells were frozen, ~1/5 was lysed for analysis and 1500 cells were seeded onto a 10-cm dish containing 1 million mouse embryonic fibroblasts (MEFs) and cultured in StemFlex media for clonal expansion. Media was replaced every other day for 7-10 days until colonies were well visible. Colonies were then picked, split in several pieces and half was transferred into a well of two matched Matrigel coated 96-well plates for maintenance and analysis, respectively.

#### v. Analysis of rs4702 genomic locus

The rs4702 region was amplified using AmpliTaq Gold™ 360 Master Mix(ThermoFisher, # 4398901) and primers against the rs4702 region (f: GGAATAGTTGAGCCCCAAGTCC, r: TGACTTGGGCCCACATCCAG). PCR conditions were as follows; 95 °C for 10 min followed by 35 cycles (95 °C for 30 s, 60 °C for 30 s, 72 °C for 60 s) and 72 °C for 7 min. The restriction enzymes SfaN1 (NEB; cuts specifically rs4702 G) and Bsr1 (NEB; cuts specifically rs4702 A) were used for further analysis of the PCR product of the bulk edit and the picked clones. After identification of potential successfully edited clones, the sequence was confirmed by Sanger using the forward primer.

#### vi. hiPSC SNP editing efficiency evaluation by ABEmax system

For testing bulk DNA base editing efficiency, we used a customized pβactin-ABEmax-puro vector system for which we cloned dCas9-adenine base editor part from ABEmax vector (Addgene #112095) ^75^ into pSpCas9(BB)-2A-Puro (PX459) V2.0 vector (Addgene# 62988) ^67^. We edited a common SNP rs7148456 that showed allele-specific open chromatin in *BAG5* ^55^ in two hiPSC lines: CW70372 (low PRS) and CW30525 (high PRS). Cells for editing were plated on 4-well dishes (Thermo Scientific) coated with Matrigel (Corning). iPSCs were cultured with mTeSR plus (StemCell) and passaged with ReLeSR (StemCell). All cells were incubated, maintained and cultured at 37°C with 5% CO_2_. Once hiPSCs reached ~70% confluency they were treated with 1.5 μl Lipofectamine STEM (Thermo Fisher Scientific) according to manufacturer’s protocols and 750 ng ABEmax BAG5 targeting plasmid. 24 hours post transfection, cells were treated with 0.5 μg/ml Puromycin (InvivoGen). 48 hours post transfection, cells were treated with 0.25 μg/ml Puromycin. 72 hours post transfection, cells were washed with DPBS (Thermo Fisher Scientific) and detached from plates with Accutase (Stem Cell). QuickEx (Lucigen) was used for rapidly extracting genomic DNAs from the transfected cells according to manufacturer’s protocol. The extracted DNAs were used as template for PCR, followed by Sanger sequencing. Primers used for genomic DNA amplification: BAG5_rs7148456_onTar_L: TTCCCCTCCCCACCCTTTTA, BAG5_rs7148456_onTar_R: CGCTCAGACCTAGTCGGGA.

### Molecular analysis

#### i. Real time-quantitative PCR

Cell were harvested with Trizol and total RNA extraction was carried out following the manufacturer’s instructions. Quantitative transcript analysis was performed using a QuantStudio 7 Flex Real-Time PCR System with the Power SYBR Green RNA-to-Ct Real-Time qPCR Kit (all ThermoFisher). Total RNA template (25 ng per reaction) was added to the PCR mix, including primers. qPCR conditions were as follows; 48C for 15 min, 95C for 10 min followed by 45 cycles (95C for 15 s, 60C for 60 s). All qPCR data is collected from at least 3 independent biological replicates of one experiment. Data analyses were performed using GraphPad PRISM 8 software.

Primers were used as follows: 18S (f: ACACGGACAGGATTGACAGA, r: *NANOG* (f: CAGCTGTGTGTACTCAATGATAGATTTC, r: GGACATCTAAGGGCATCACAG); GGAGAATTTGGCTGGAACTGCATG); *PAX6 (*f: GGAGTGAATCAGCTCGGTGG, r: GGTCTGCCCGTTCAACATCC); *LMX1A* (f: TCCAGGTGTGGTTCCAAAAC, r: GGTTCATGATTCCTTCCATCCC); *HAND1 (*f: AAAGGCTCAGGACCCAAGAAG, r: TGATCTTGGAGAGCTTGGTGTC); *HOPX* (f: CGAGGAGGAGACCCAGAAATG, r: GACGGATCTGCACTCTGAGG); *FOXP2 (*f: CAGTCACCCCGATTACCCAG, r: GGGGCAATTTCTGATGACATGG); *SOX17 (*f: GAACGCTTTCATGGTGTGGG, r: CACGACTTGCCCAGCATCT); *GAD2* (ThermoFisher, Hs00609534_m1) and *vGAT* (ThermoFisher, Hs00369773_m1).

#### ii. Immunostaining and microscopy

Cells were washed with ice-cold PBS and fixed with 4% PFA/sucrose PBS solution at pH 7.4 for 20 mins, room temperature. Then, fixative solution was replaced with permeabilizing solution (0.1% Triton-X PBS) for 20 mins. After washed with PBS, cells were incubated with blocking solution (5% donkey serum in PBS) for 1 hour, at room temperature. The blocking solution was aspirated and replaced with the same solution with primary antibodies (MAP2-Ck, Abcam ab5392, 1:500; Nanog-Gt, R&D AF1997, 1:200; TRA-1-60 IgM-Ms, Millipore MAB4360, 1:100; GFAP-Ck Aves AB_2313547, 1:1000; MAP2AB-Ms, Sigma M1406, 1:500; synapsin1-Ms Synaptic systems, 1:500; TH-Rb, Pel-Freez P40101, 1:1000; S100beta-Ms, Sigma S2532, 1:1000; HuNu, Sigma MAB1281B, 1:200; MAP2, Synaptic Systems 188 003, 188 011, 1:700; and GABA, 1:1000, Sigma A2052) overnight at 4°C. Cells were then incubated with secondary antibodies (Alexa 488 anti-Rabbit, Dk, Jackson immunoResearch, 711-545-152, 1:500; Alexa 568 anti-Chicken, Gt, ThermoFisher, A-11041, 1:500; Alexa 647 anti-mouse, Dk, Jackson immunoResearch, 715-605-150, 1:500), prepared in blocking solution, for two hours at room temperature, followed by DAPI staining for 3-4 min and PBS-washing 3 times.

Cells were imaged with a Zeiss LSM 780, Nikon C2 confocal microscope or a ThermoFisher HCS CX7 microscope. The images were analyzed using ImageJ, except astrocytes, which were quantified using the CellProfiler software. Data points represent 10-33 images from 2 biological replicates.

## DATA AVAILABILITY

All source donor hiPSCs are available from the CIRM repository (https://www.cirm.ca.gov/researchers/ipsc-repository). Genotype data will be made publicly available prior to publication.

## ACKNOWLEDGEMENTS

This work was supported by a supplement to National Institute of Health (NIH) grant R01MH113215 (D.A.), R56 MH101454 (K.J.B.), R01MH106575 (J.D.) and R01MH116281 (J.D.), and U01 MH115727 (SM and KE).

Special thanks to Dr. David Panchision at NIMH for program guidance on this collaborative project. All hiPSC lines were obtained from the CIRM hPSC Repository funded by the California Institute of Regenerative Medicine (CIRM). Portions of Figures 1 and 5 were created with BioRender.com

## AUTHOR CONTRIBUTIONS

K.J.B, J.D. and D.A. conceptualized this collaborative approach. K.J.B, J.D., D.A, K.R., H.Z., S.A. contributed to experimental design and wrote the manuscript. S.G., K.E. and S.M. identified the high and low PRS hiPSC lines in the CIRM hPSC repository. K.R., H.Z., T.P., S.W., D.D., S.A, conducted all pluripotency experiments, transfections, neuron production, neuronal differentiation and CRISPR-editing experiments. D.A. conducted all genomic and CNV analyses on the hiPSC lines.

